# Neuronal age drives β-sheet accumulation and, together with ApoE4, enhances synaptic Aβ localization in APP^NL-F^ neurons

**DOI:** 10.64898/2026.01.10.697647

**Authors:** Sabine C Konings, Emma Nyberg, Isak Martinsson, Radhika Thakore, Gunnar K. Gouras, Oxana Klementieva

## Abstract

Amyloid-β (Aβ) accumulation and aggregation are defining features of Alzheimer’s disease (AD), but the earliest cellular events driving these processes remain poorly understood. Here, we investigate Aβ dynamics in primary neurons derived from APP^NL-F^ knock-in mice, which express human APP and Aβ at endogenous (non-overexpressed) levels. Using correlative optical photothermal infrared (OPTIR) spectromicroscopy and immunofluorescence, we show that physiological APP^NL-F^ expression is sufficient to drive intraneuronal Aβ accumulation at synapses. Modelling neuronal aging through prolonged culture reveals a marked increase in synaptic Aβ burden accompanied by β-sheet–rich structural transitions Supplementation with astrocyte-derived apolipoprotein E (ApoE) isoforms demonstrates that ApoE4 markedly increases synaptic Aβ accumulation while reducing β-sheet content relative to untreated cultures, suggesting a potential shift toward less fibrillar species. Together, these findings establish APP^NL-F^ neurons as a physiologically relevant system for dissecting early Aβ pathology and show that both Aβ quantity (accumulation) and quality (structural maturation) are regulated by aging and ApoE genotype. This work provides mechanistic insight into the earliest molecular events that may underlie synaptic vulnerability in AD.

## Introduction

Alzheimer’s disease (AD) is an age-dependent neurodegenerative disorder characterized by abnormal protein aggregation, including amyloid-β (Aβ), which accumulates in the brain and ultimately forms amyloid plaques.^1,2^ Multiple Aβ assemblies have been identified—soluble oligomers, protofibrils, and mature fibrils—each exhibiting distinct structural and functional properties that may differentially contribute to neurodegeneration.^3,4^ In the early cellular phase of AD, before plaque formation, Aβ accumulates intracellularly in neurons.^5^ Converging biophysical, cellular, and histological evidence indicates that low-molecular-weight Aβ oligomers—small assemblies of misfolded monomers—are particularly neurotoxic and strongly implicated in synaptic dysfunction.^6–9^

Intraneuronal Aβ accumulation is especially prominent at synaptic terminals,^10^ which are among the earliest cellular sites affected in AD. Synapse loss correlates strongly with cognitive decline in patients.^11,12^ Studies of synaptosomes isolated from AD brains^13,14^ have revealed that synaptic Aβ is predominantly oligomeric. Moreover, synaptic Aβ is labeled by both prefibrillar and fibrillar Aβ antibodies, indicating structural heterogeneity of Aβ species at synapses.^15^ Experimental application of soluble Aβ oligomers induces synaptic deficits, including impaired synaptic plasticity, abnormal spine morphology, and eventual synapse loss.^7,9,16^ While intracellular synaptic Aβ accumulation has been described in primary neurons derived from transgenic APP-overexpressing models,^17,18^ it has not yet been demonstrated in primary neurons expressing human APP at physiological levels.

Although recent anti-Aβ antibody therapies have brought the field closer to slowing AD progression, ^19^ effective treatments remain limited,^20^ and many therapeutic strategies that show promise in transgenic mouse models fail to translate to human patients.^21^ A major limitation of conventional transgenic AD models is the artificial overproduction of APP and Aβ, which alters aggregation. To overcome this, Saito et al., (2014) developed APP knock-in (APP-KI) mouse models in which APP is expressed under the endogenous mouse promoter, thereby producing Aβ at more physiological levels.^22^ Among these, APP^NL-F^ mice carry the Swedish (NL) and Iberian (F) mutations, resulting in increased Aβ(1–42) production, accumulation, and amyloid plaque formation, and recapitulating AD-like synaptic and cognitive phenotypes.^22,23^

Despite extensive *in vivo* characterization of APP^NL-F^ mice,^22–25^ relatively little is known about intraneuronal Aβ accumulation and aggregation in APP^NL-F^ primary neurons. Primary neurons are widely used to study cellular mechanisms and may provide a more physiologically relevant AD model than earlier systems. To date, only a few studies have addressed this model: Dentoni et al. (2022) reported an elevated Aβ42/Aβ40 ratio together with mitochondrial and synaptic alterations,^25^.while Yu et al. (2025) observed increased APP C-terminal fragment accumulation in presynaptic terminals.^24^ However, how intraneuronal Aβ evolves over time in culture—a proxy for neuronal aging—and how apolipoprotein E (ApoE), the strongest common genetic risk factor for late-onset AD, modulates Aβ accumulation and aggregation in this context remain unclear.

ApoE4 has been associated with increased Aβ aggregation and enhanced deposition in the brain parenchyma.^26–28^. Lipid-associated ApoE showed higher binding of ApoE3 than ApoE4 to monomeric Aβ40 and Aβ42.^29^ This observation was complemented with later *in vitro* studies reporting preferential binding of ApoE4 to intermediate Aβ aggregates, where ApoE4 forms more complexes than ApoE2 or ApoE3.^30^ It has been also shown that ApoE lipidation and pH can affect interaction^31^ with Aβ. ApoE4 also impairs Aβ clearance,^26,32^ increases intraneuronal and synaptic Aβ accumulation,^33–36^ and reduced neuronal Aβ trafficking to lysosomes.^37,38^ Moreover, ApoE isoforms differentially regulate Aβ uptake, recycling, and degradation, with ApoE4 promoting Aβ accumulation in vulnerable neuronal compartments.^14,35,38^ Together these observations indicate that ApoE–Aβ interactions are highly context dependent, with ApoE lipidation state, pH and potentially other factors strongly modulating binding behavior.

However, whether ApoE4 influences intracellular Aβ aggregation and β-sheet structural transitions in APP-KI neurons remains unknown. To assess structural changes, we used infrared microspectroscopy a powerful label-free approach to interrogate protein secondary structure *in situ*, as the Amide I band (1600–1700 cm⁻¹) is highly sensitive to β-sheet formation—an essential molecular signature of Aβ aggregation.^39^ OPTIR overcomes the diffraction limits of traditional FTIR or QCL-IR microscopy by detecting photothermal modulation of a visible probe laser, thereby achieving submicron spatial resolution.^40^ OPTIR can resolve major and minor β-sheet components characteristic of fibrillar structures without requiring chemical labels, providing direct, spatially localized information on β-sheet–associated structural features. Importantly, OPTIR provides qualitative information on local protein secondary structure; β-sheet–associated spectral intensity reflects structural organization at the measurement site rather than total β-sheet content. When combined with immunofluorescence, OPTIR enables specific assignment of β-sheet–rich spectral signatures to amyloids, thereby linking molecular identity to structural fingerprints within single cells.^41^

In this study, we investigated Aβ accumulation and β-sheet formation in hAPP^NL-F^ primary neurons at the subcellular level, focusing on how key AD risk factors—neuronal aging and ApoE genotype—shape intraneuronal Aβ. APP^NL-F^ primary neurons were cultured for 15 days *in vitro* (DIV) to generate mature neurons and for 25 DIV to model neuronal aging. Structural alterations associated with β-sheet formation and lipid composition were examined usings the Swedish (NL) and Iberian (F) Super–Resolution infrared microspectroscopy,^42^ while fluorescence microscopy and biochemical analyses were used to evaluate synapse morphology and synaptic Aβ localization. Finally, by supplementing neurons with astrocyte-derived human ApoE3 or ApoE4, we assessed isoform-dependent effects on Aβ accumulation, aggregation, and synaptic morphology in APP^NL-F^ neurons.

## Results

### APP^NL-F^ primary neurons contain β-sheet signatures consistent with amyloid assemblies

Primary neurons derived from transgenic AD mouse models have been widely used to study intracellular Aβ accumulation.^17,18,36^ In contrast, primary neurons derived from APP knock-in mice, expressing more physiological levels of human Aβ, remain largely unexplored. We therefore used APP^NL-F^ knock-in mice, in which humanized APP is expressed from the endogenous promoter and carries the Swedish (NL) and Iberian (F) AD mutations, resulting in more physiological Aβ production without APP overexpression.^22^ To confirm the presence of human Aβ in mature APP^NL-F^ primary neurons, we used two antibodies: 6E10, which recognizes the Aβ domain of human APP, and 82E1, which detects the free Aβ N-terminus. Wild-type neurons were used as baseline control to account for nonspecific antibody binding. Both antibodies showed clear signal in neuronal cell bodies (**Supplementary Figure 1A–B**) as well as along neurites (**Supplementary Figure 1C–D**) of APP^NL-F^ neurons. In contrast, WT murine neurons showed only slight background labelling with human-specific antibody 6E10 and markedly less labelling with N-terminal antibody 82E1.

While neuronal aging cannot be directly modeled *in vitro*, prolonged time in culture induces progressive changes that recapitulate age-associated signatures.^43–46^ To examine the formation of β-sheet structures in neurons before and during this aging trajectory, we analyzed two culture time points: 15 days *in vitro* (DIV), representing mature neurons, and 25 DIV, representing aged neurons. To directly assess protein secondary structure in intact neurons, we used optical photothermal infrared spectroscopy (OPTIR).^40^ OPTIR resolves vibrational signatures at ∼500 nm spatial resolution, thus enabling detection of β-sheet signatures characteristic of amyloid assemblies without chemical labeling at subcellular level.^42^ In parallel, OPTIR captures lipid-associated ester vibrations (∼1740 cm⁻¹) indicative of lipid enrichment or oxidation,^47^ allowing simultaneous readout of protein misfolding and membrane chemistry within single neurons (**Table 1**). We observed that a subset of spectra acquired from APP^NL-F^ neurons at 15 and 25 DIV exhibited an increase at 1628 cm⁻¹, corresponding to elevated β-sheet structural content characteristic of amyloid fibrils.^39^ Importantly, this feature was not present in all spectra acquired from a given neuron, therefore we pooled all spectra from APP^NL-F^ neurons and classified them into two groups: APP^NL-F^ (β-sheet–positive) and APP^NL-F^ (non-β-sheet) cohorts (**Figure 1B,F)**. Spectral analysis revealed that (i) spectra from wild-type control neurons did not show an increase in β-sheet content over time (**Figure 1D**, grey bars); (ii) non-β-sheet spectra from APP^NL-F^ neurons did not exhibit changes in intensity at the 1628 cm⁻¹ (**Figure 1D**, magenta bars); and (iii) β-sheet–positive spectra were already detectable at 15 DIV in APP^NL-F^ neurons (p<0.0001) as compared to wild type control and significantly increased (p<0.001) with time in culture (15 vs 25 DIV) (**Figure 1D**, red bars). Because the amount of Aβ present within an individual measurement site (∼500 nm spot size) cannot be quantitatively determined, the analysis was restricted to qualitative assessment of the local protein conformational state, specifically the presence of β-sheet–rich structures relative to other proteins within the sampled volume. Alongside β-sheet emergence, the same spectra revealed increased intensity at ∼1738 cm⁻¹, corresponding to ester carbonyl vibrations—a signature associated with lipid enrichment or oxidation.^47^ This lipid-associated band increased with DIV in both wild-type and non-amyloid APP^NL-F^ neurons (**Figure 1H**). Notably, in APP^NL-F^ β-sheet positive spectra this age-dependent lipid increase was not observed. Together, these data indicate that β-sheet formation in APP^NL-F^ neurons is associated with a distinct lipid profile that differs from the DIV-dependent lipid changes observed in wild-type neurons and the β-sheet negative cluster of APP^NL-F^ neurons.

**Table 1.** OPTIR spectral assignments for protein secondary structures and lipids.

| Molecular Class | Band (cm<sup>-1</sup>) | Assignment | Interpretation / Relevance |
| --- | --- | --- | --- |
| Proteins | 1625–1638 | β-sheet, major component | Aggregated / amyloid-like structures |
| | 1648–1660 | α-helix | Native protein secondary structure |
|  | 1660–1680 | Turns | Protein structural heterogeneity |
| | 1680–1695 | $\beta'$-sheet, minor component | Aggregation, oligomeric/fibrillar amyloids |
| Lipids | 1738 | -CO=O | Ester group |

**Figure 1.**
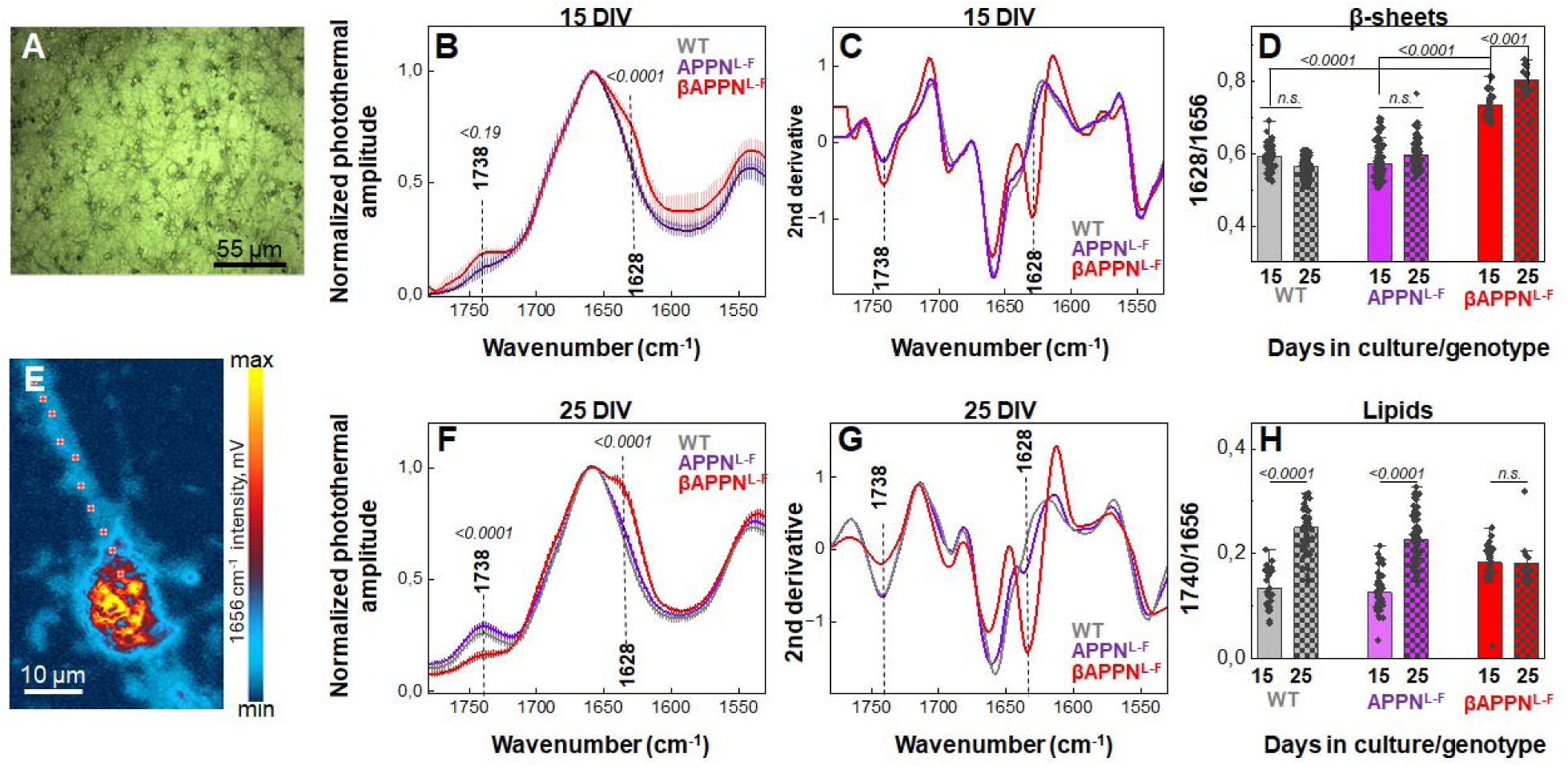
β-sheet formation in APP^NL-F^ neurons over time. **A.** Bright-field image of neurons grown on CaF2, objective 10x. **B, F**. Averaged and normalized OPTIR infrared spectra recorded from neurons grown 15 days (**B**) and 25 days (**F**) in culture. The spectral change corresponding to β-sheet structures is indicated by a vertical dashed line, the band corresponding to lipids (COO-) is indicated by a blue dashed line. **C, G**. Averaged and normalized second derivatives derived from the OPTIR spectra shown in B and F correspondingly. Vertical dashed lines indicate band positions that were analyzed. **E**. OPTIR image of a neuron at 1656 cm^-1^. Measurement positions are indicated by red dots. Red crosses show how spectral locations acquired from the neurites. Coloured bar shows the intensity of OPTIR: yellow maximum, cell body, blue – neurites, bark bule background. **D, H**. Statistical comparisons were done using OriginPro^R^ by one-way ANOVA followed by Tukey’s test, with the significance level set to 0.05, the bar shows mean ± SD. Spectra were acquired from 20–50 cells per genotype. Low-quality spectra were excluded prior to analysis. The remaining spectra were clustered, and only β-sheet–positive clusters were retained for quantification

### Aβ localizes to synapses in APP-KI^NL-F^ neurons, which is more apparent in older neurons

While OPTIR directly resolves protein secondary structures at submicron resolution, definitive protein attribution requires immunofluorescence. Combining OPTIR with antibody labeling therefore enables β-sheet spectral signatures to be assigned to Aβ, linking structural fingerprints to molecular identity. To confirm that APP^NL-F^ neurons also accumulate Aβ, we labeled mature (15 DIV) and aged (25 DIV) APP^NL-F^ neurons with the N-terminus–specific antibody 82E1 (amino acids 1–4). 82E1-positive puncta were distributed along MAP2-positive neurites, with some puncta overlapping with MAP2 and others not, consistent with localization to dendrites and synaptic regions, respectively (**Figure 2A**). This synaptic localization of Aβ was already present at 15 DIV and became more prominent at 25 DIV (**Figure 2E**, left panels; **Supplementary Figure 2**, magnifications 1–2). Despite only a modest increase in Aβ puncta number with age (+25% at 25 DIV, p = 0.0664; **Figure 2B–C**), Aβ signal intensity increased more substantially, rising by 34% from 15 to 25 DIV (p = 0.0195; **Figure 2D**). Thus, neuronal aging primarily increases the local concentration of synaptic Aβ, more than the frequency of Aβ puncta.

**Figure 2:**
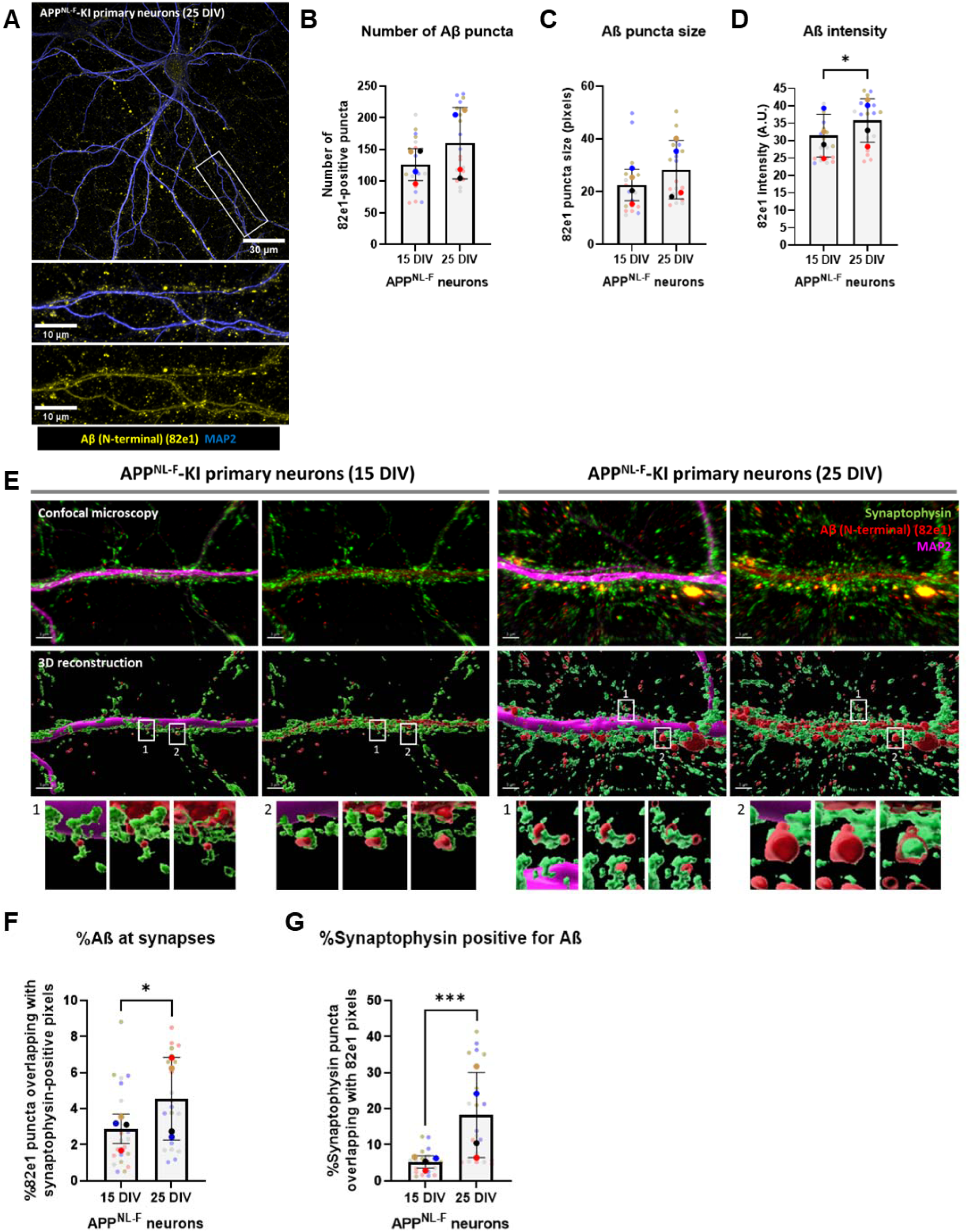
Aβ localizes to synapses in APP-KI^NL-F^ neurons, with highest amount in older neurons. **A.** Representative epifluorescence microscopy image of 25 DIV APP^NL-F^ primary neurons showing that Aβ labeling (yellow) follows a synaptic pattern along MAP2-positive dendrites (blue). Aβ was labeled using 82e1 antibody. The upper panel shows an overview of an Aβ-labeled APP^NL-F^ neuron. Scale bar in this panel represents 30 µm. The lower panel shows a higher magnification image of the white box outlined in the upper panel. Scale bar in the lower panel is 10 µm. **B – D.** Quantification of the Aβ-positive puncta number (**B**), puncta size (**C**) and puncta intensity (**D**) in APP^NL-F^ primary neurons that are mature (15 DIV; **Supplementary Figure 2**) or aged (25 DIV; Figure 2A). The larger data points reflect the number of independent embryos used (N = 4), the smaller data points indicate the number of individual neurons studied (n = 16) The statistical analyses were performed on a neuron level. **E.** Representative confocal (upper panel) and 3D reconstructed (middle panel) images of mature (15 DIV) and older (25 DIV) APP^NL-F^ primary neurons labeled for N-terminal Aβ (82e1; red), synaptic marker synaptophysin (green), and MAP2 (magenta). The lower panels show 2 examples of Aβ at synaptophysin-positive synapses per condition at a higher magnification. Scale bar is 3 µm. **F – G.** Quantification of the percentage of pre-synaptic synaptophysin pixels that are positive for Aβ (**F**) and the percentage of Aβ pixels that is colocalizing with synaptophysin-positive synapses (**G**) in mature (15 DIV) and older (25 DIV) APP^NL-F^ primary neurons. Bigger data points indicate the number of embryos (N = 4), smaller data points are reflecting individual neurons (n = 16). Data is shown as mean ± SD. *p ≤ 0.05, ***p ≤ 0.001.

To directly test whether these puncta were synaptic, we co-labeled APP^NL-F^ neurons with 82E1 and the presynaptic marker synaptophysin. Three-dimensional reconstructions of confocal z-stacks revealed clear co-localization of Aβ with synaptophysin (**Figure 2E**), confirming that the accumulated Aβ resides at synaptic terminals. At 15 DIV, co-localization was detectable but sparse. By contrast, in aged cultures, Aβ occupied synapses in two distinct modes: either as small puncta opposed to synaptophysin terminals (**Figure 2E**, 25 DIV panel 1), or as larger deposits that fully enveloped synaptophysin-positive boutons (F**igure 2E**, 25 DIV panel 2), a pattern not observed in mature (15 DIV) neurons.

Quantitatively, the proportion of Aβ overlapping with synaptophysin increased by 109%, from 2.05% at 15 DIV to 4.27% at 25 DIV (p = 0.0283; **Figure 2F**). Similarly, the fraction of synaptophysin-positive terminals containing Aβ increased by 154%, from 15 to 25 DIV (p = 0.0002; **Figure 2G**).

Together, these data show that Aβ accumulates at synapses and that aging significantly increases synaptic Aβ burden. Neuronal aging particularly drives increased Aβ concentration within individual boutons.

### Time in culture influences synaptic protein levels in neurons independently of the presence of human APP/Aβ

Synaptic disruption is among the earliest cellular manifestations of AD, emerging before neuronal loss and correlating more strongly with cognitive decline than plaque burden.^12,48^ We therefore hypothesized that subtle synaptic alterations would arise in primary neurons as Aβ accumulates and begins to aggregate, and that these changes would progress with neuronal age. To test this, we performed SDS–PAGE followed by Western blot analysis on lysates from wild-type and APP^NL-F^ primary neurons cultured for 15 or 25 days *in vitro* (DIV).

Using pre- and postsynaptic markers, we observed robust age-dependent increases in presynaptic proteins, including Synaptophysin, Rab3a, and Bassoon (**Figure 3A–D**; p = 0.0002, p = 0.0003, and p = 0.0012, respectively). Importantly, similar changes were observed for both wild-type and APP^NL-F^ neurons.

**Figure 3:**
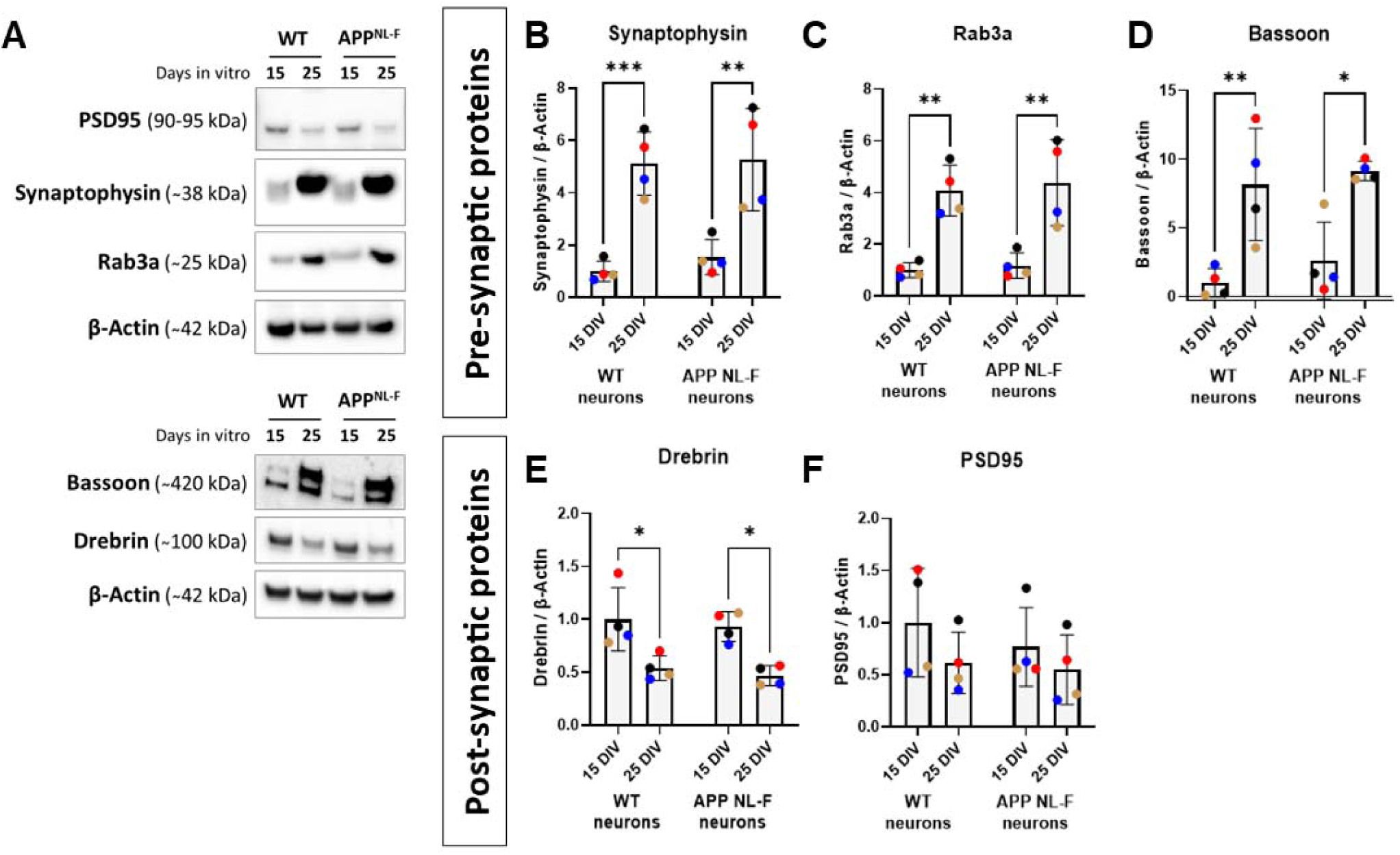
Pre-synaptic protein levels are increased in older (25 DIV) primary neurons, independent of the presence of human APP. **A.** Representative western blot membranes showing the protein bands of various pre-synaptic (synaptophysin, Rab3a and Bassoon) and post-synaptic proteins (PSD95 and Drebrin) in neuron lysate obtained from wild-type and APP^NL-F^ primary neurons (15 and 25 days *in vitro* (DIV)). **B-F.** Quantification of the protein levels of pre-synaptic synaptophysin (**B**) and Rab3a (**C**), synaptic cleft protein Bassoon (**D**), and post-synaptic Drebrin (**E**) and PSD95 (**F**) in mature (15 DIV) and older (25 DIV) wild-type and APP^NL-F^ primary neurons. One data point is showing a single embryo (N = 4). Data is shown as mean ± SD. *p ≤ 0.05, **p ≤ 0.01, ***p ≤ 0.001.

In contrast, the levels of postsynaptic proteins decline with age. Drebrin levels decreased across both genotypes (**Figure 3A–E**; p = 0.0003), and PSD95 exhibited a similar downward trend (**Figure 3A–F**; p = 0.0268), although genotype-specific comparisons did not reach statistical significance (WT: p = 0.0768; APP^NL-F^: p = 0.3367). The parallel age-associated shifts in wild-type and APP^NL-F^ neurons indicate that these changes reflect a general aging process rather than a direct consequence of human APP expression.

### Altered synaptic morphology in mature APP^NL-F^ primary neurons

Synaptic disruption is an early hallmark of AD, that is thought to emerge prior to neuronal loss, and correlates more strongly with cognitive decline than plaque burden. We next wondered if structural abnormalities may precede measurable changes in synaptic protein abundance and therefore examined whether APP^NL-F^ neurons exhibit synaptic remodeling.

Using confocal microscopy, we visualized the presynaptic marker synaptophysin together with Bassoon, a core scaffold of the active zone. APP^NL-F^ neurons displayed a distinct synaptic morphology relative to wild-type cultures (**Figure 4B**). Synaptophysin-positive puncta frequently appeared elongated and enlarged rather than rounded (**Figure 4 A,B**), consistent with a structural remodeling phenotype rather than synapse loss.

**Figure 4:**
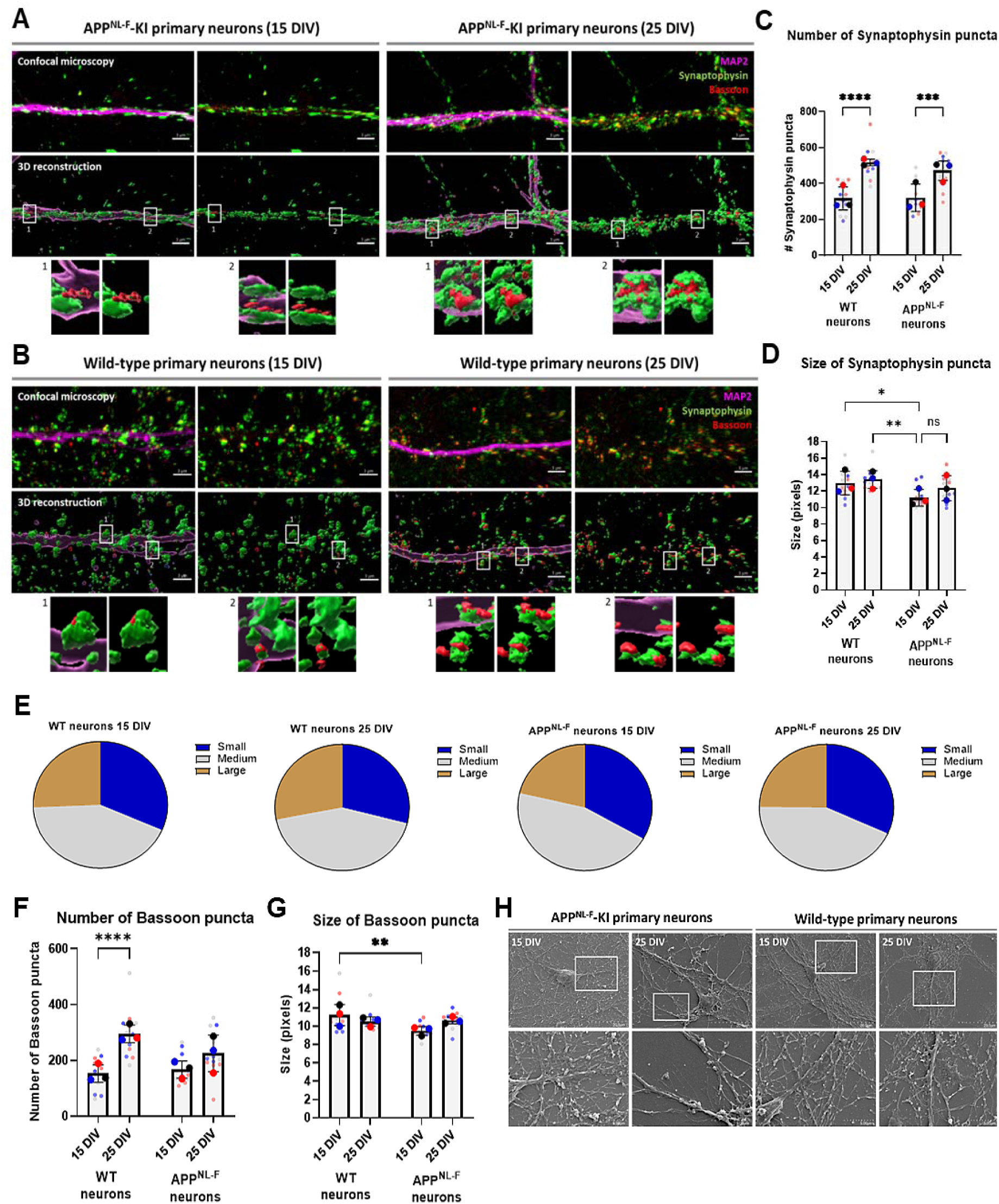
Synaptic morphology is altered in APP^NL-F^ primary neurons, particularly in mature cells. A – B. Confocal (upper panel) and 3D-reconstructed (middle panel) images of mature (15 DIV) and older (25 DIV) APP^NL-F^ (A) and wild-type (B) primary neurons showing synaptic puncta for synaptophysin (green) and bassoon (red) along MAP2-labeled dendrites (magenta). The lower panels show higher magnification images of synaptophysin-bassoon-interaction at the synapses (two examples for each condition were shown). Note: Synaptic labeling in mature (15 DIV) APP^NL-F^ neurons show more a linear structure, while the other conditions have more round-shaped synaptic puncta. Scale bar is 3 µm. C – D. Quantification of the number (C) and size (D) of the synaptophysin-positive puncta in mature (15 DIV) and older (25 DIV) APP^NL-F^ and wild-type primary neurons. Bigger points reflect embryos (N = 3), smaller points reflect neurons (n = 12). Statistics were done on a neuron level. E. Pie charts of the size distribution of the synaptophysin puncta in wild-type (15 and 25 DIV) and APP^NL-F^ (15 and 25 DIV) primary neurons. Small puncta (blue) are all puncta smaller than the average first quartile, large puncta (gold) are all puncta larger than the average third quartile, medium sized puncta (blue) are all puncta in between the average first and the third quartile. F – G. Quantification of the bassoon-positive synapse puncta number (F) and puncta size (G) in wild-type (15 and 25 DIV) and APP^NL-F^ (15 and 25 DIV) primary neurons. Larger data points indicate embryos (N = 3), smaller data points indicate individual analyzed neurons (n = 12). **H**. Representative image of APP^NL-F^ and wild-type primary neuron cultures taken by a scanning electron microscope. These images show synaptic morphology in mature (15 DIV) APP^NL-F^ primary neurons (as shown by less protrusions and branching from the neurite) compared to the other conditions. Data is shown as mean ± SD. *p ≤ 0.05, **p ≤ 0.01, ***p ≤ 0.001, ****p ≤ 0.0001.

Quantitative analysis revealed a robust age-dependent increase in synaptophysin puncta number in both genotypes (**Figure 4C**; wild-type: +63.4%, p = 0.0001; APP^NL-F^ +46.9%, p = 0.0006), mirroring the elevated synaptophysin protein levels observed by western blot (**Figure 3B**). Notably, synaptophysin puncta were significantly smaller in 15 DIV APP^NL-F^ neurons compared to 15 and 25 DIV wild-type neurons (**Figure 4D**; −13.8%, p = 0.0202; −16.6%, p = 0.0025), whereas overall puncta intensity remained unchanged (**Supplementary Figure 3D**). Unlike 15 DIV, 25 DIV APP^NL-F^ neurons did not show a difference in size of synaptophysin puncta from wild-type neurons (**Figure 4D**). Size-distribution analysis further showed that 15 DIV APP^NL-F^ neurons contained fewer large (> third quartile) synaptophysin-positive synaptic terminals (**Figure 4E**), indicating a shift toward structurally reduced presynaptic compartments. This change was significant relative to 25 DIV wild-type neurons (p = 0.0013; **Supplementary Figure 3C**), while the proportion of small and medium synapses remained stable (**Supplementary Figure 3A-B**).

Bassoon puncta showed a similar but partially genotype-dependent trend. Wild-type neurons displayed a pronounced age-driven increase (**Figure 4F**; +93.1%, p = 0.0001), whereas APP^NL-F^ neurons exhibited only a modest, non-significant change (**Figure 4F**; +34.8%, p = 0.1008). Consistent with synaptophysin, Bassoon puncta were significantly smaller in 15 DIV APP^NL-F^ neurons compared to 15 DIV wild-type (**Figure 4G**; −15.0%, p = 0.0064) but no difference was detected at 25 DIV. Importantly, Bassoon puncta intensity also was less in 15 DIV APP^NL-F^ cultures (**Supplementary Figure 3E**; −41.7%, p = 0.0124), suggesting change presynaptic active zone assembly rather than an increase in synapse number.

To independently assess synaptic morphology, we performed scanning electron microscopy (SEM). While fluorescent puncta-based analyses primarily revealed early synaptic alterations in APP^NL-F^ neurons at 15 DIV, SEM showed pronounced structural alterations at 25 DIV. APP^NL-F^ neurons at 25 DIV exhibited elongated protrusions (**Figure 4H**), in contrast to the morphology in wild-type neurons and in 15 DIV APP^NL-F^ cultures (**Figure 4H**). These ultrastructural changes mirror the elongation phenotype detected by immunofluorescence, reinforcing the notion of synaptic structural remodeling rather than terminal loss.

### Effect of ApoE on β-sheet content

ApoE4 is the strongest genetic risk factor for late-onset AD.^49,50^ In human post-mortem AD brain tissue, ApoE co-localizes with oligomeric Aβ at synaptic terminals,^14,35^ suggesting that ApoE–Aβ interactions occur precisely within the compartments most vulnerable to early pathology. Studying this interaction in experimental models is therefore critical for understanding how ApoE shapes the earliest stages of Aβ accumulation and aggregation.

To address ApoE isoform–dependent effects on Aβ secondary structure, we combined OPTIR and confocal imaging on APP^NL-F^ primary neurons. Because the APP^NL-F^ mouse retains endogenous mouse ApoE, we introduced human ApoEs by exposing neurons to astrocyte-conditioned media from primary astrocyte derived humanized ApoE3-KI and ApoE4-KI mice. Both wild-type and APP^NL-F^ neurons were incubated for 8 hours with ApoE3- or ApoE4-conditioned media, which enabled us to assess isoform-specific effects on β-sheet structural signatures in the presence of astrocyte-derived human ApoE (**Figure 5A**). Conditioned media from ApoE-knock out astrocytes served as the control condition. ApoE treatment resulted in a reduction of the major β-sheet component at 1628 cm⁻¹, with the strongest decrease observed in the ApoE4 condition (**Figure 5B**). To resolve the secondary-structure subcomponents of the Amide I band, we performed second-derivative analysis,^51^ identifying peaks at 1656 cm⁻¹ (α-/unordered), 1628 cm⁻¹ (major β-sheet), and 1690 cm⁻¹ (minor antiparallel β′-sheet) (**Supplementary Figure 5**). The fitted Amide I spectra (**Figure 5E–G**) revealed distinct secondary-structure fingerprints across the different ApoE conditions. Two major peaks were identified, corresponding to major and minor β-sheet components, together with contributions from α-helical and turn structures. Notably, treatment with ApoE4- and ApoE3-conditioned media resulted in a decrease in the fitted component corresponding to the major β-sheet structure, manifested as a reduced peak centered at ∼1628 cm⁻¹, indicating structural rearrangements induced by ApoE treatment (**Figure 5F,G**). We speculate that the observed reduction in β-sheet content in APP^NL-F^ neurons after ApoE treatment, particularly under ApoE4 exposure, may cause a shift in Aβ structural organization. This shift is consistent with reports that ApoE4 binds intermediate Aβ species and enhances oligomer retention rather than promoting fibril formation. ^31,37^

**Figure 5.**
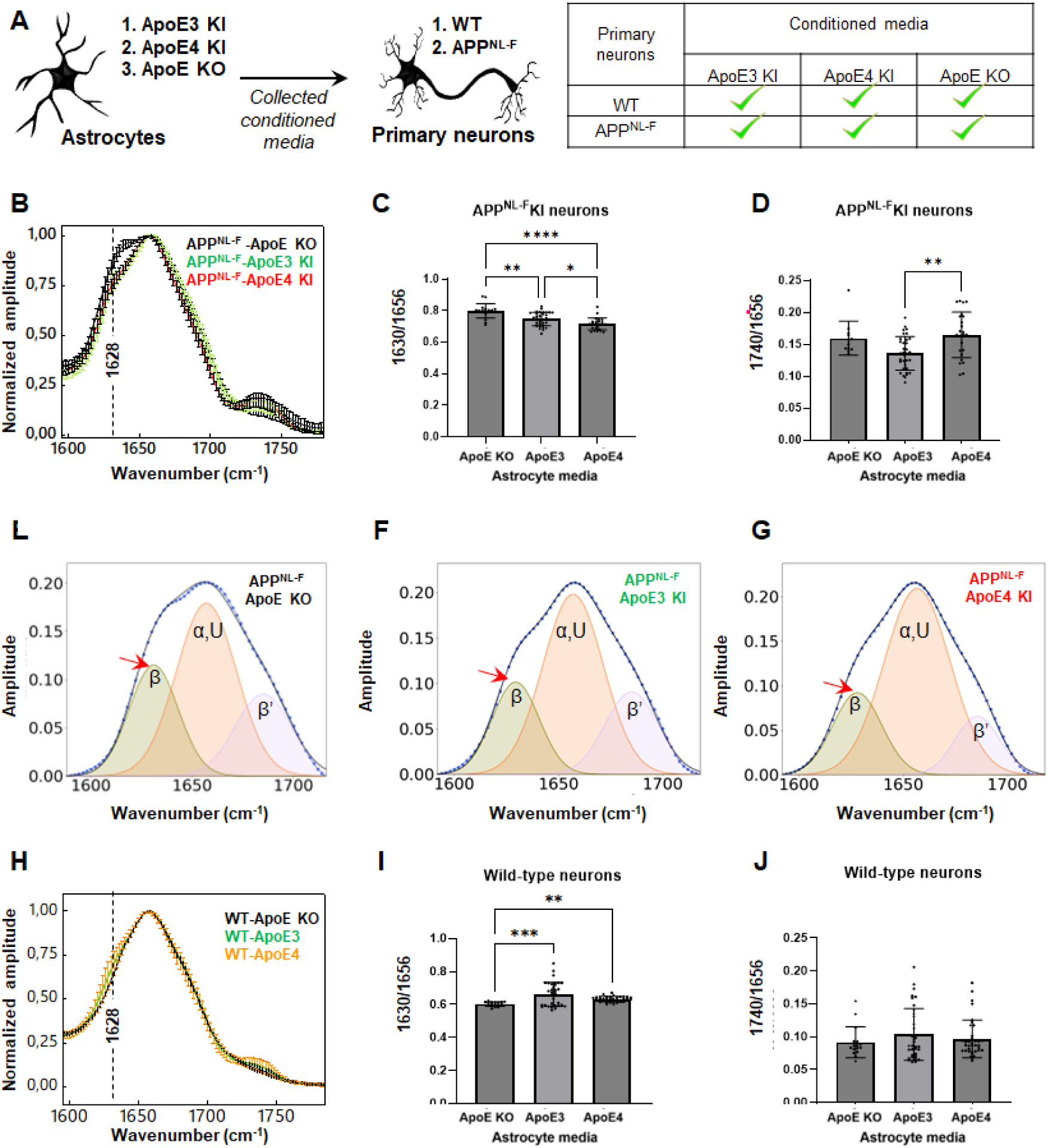
Effect of ApoE on amyloid formation in APP^NL-F^ neurons. **A**. Schematic overview of our approach to add human astrocyte-derived ApoE3 and ApoE4 to aged APP^NL-F^ primary neuron cultures (25 DIV). In short, human ApoE3 and ApoE4 (or ApoE KO) was derived from primary mouse astrocytes obtained from the humanized ApoE3-KI, ApoE4-KI or ApoE KO mice. The astrocyte conditioned media from these mice was added as a treatment to aged APP^NL-F^ primary neurons (25 DIV) for 8 hours. After 8 hour incubation, the neurons were fixed and used for further analysis. **B.** Averaged and normalized OPTIR infrared spectra recorded from APP^NL-F^ neurons treated with ApoE3 and ApoE4. at 25 days in culture. The spectral feature corresponding to β-sheet structures is indicated by a red dashed line, the bar shows mean ± SD (corresponding 2^nd^ derivatives are shown in **Supplementary** Figure 5A). **C, D.** Statistical comparisons of β-sheets and lipids were done by one-way ANOVA followed by Tukey’s test, with the significance level set to 0.5, the bar shows mean ± SD. We examined 3 embryos per genotype, 20 to 50 cells per embryo. **E-G**. Spectral deconvolution of the amide I IR spectra shows the individual secondary structure components. The raw spectrum is shown as a black line, and the fitted curve as a blue dotted line. The main β-sheet component (orange, band centered around 1630 cm⁻¹), the mixed α-helix/unordered structures (purple, band centered around 1656 cm⁻¹), and the antiparallel β′-sheet component (green, band centered around 1690 cm⁻¹) are indicated. The arrow highlights the increased β-sheet component observed in APP^Nl-F^ neurons. **H.** Averaged and normalized OPTIR infrared spectra recorded from wild-type neurons grown 25 days in culture and treated with ApoE3 and ApoE4. The spectral feature corresponding to β-sheet structures is indicated by a red dashed line, the bar shows mean ± SD. Corresponding 2^nd^ derivatives are shown in **Supplementary** Figure 5B. **J,H.** Statistical comparisons of β-sheets and lipids were done by one-way ANOVA followed by Tukey’s test, with the significance level set to 0.05, the bar shows mean ± SD. We examined 3 embryos per genotype, 20 to 50 cells per embryo.

Since ApoE is directly involved in lipid transport and membrane metabolism, we next examined whether ApoE exposure was associated with changes in lipid-associated spectral features by quantifying the ester-associated band at 1740 cm⁻¹. Lipid content differed significantly between ApoE isoforms, with ApoE4-treated neurons exhibiting higher lipid spectra intensity than ApoE3-treated neurons (**Figure 5D**). Such increased lipid content may reflect lipid accumulation or altered membrane remodeling, consistent with the impaired neuronal lipid efflux previously reported with ApoE4 ^52,53^

Finally, treating wild-type neurons with the same astrocyte-conditioned media resulted in only a small increase in β-sheet content with ApoE3 or ApoE4 (**Figure 5H–J**), which is a much weaker effect than in APP^NL-F^ neurons. This indicates that the structural changes detected by OPTIR are primarily linked to interactions between ApoE isoforms and human Aβ, although contributions from other aggregation-prone proteins cannot be excluded.

### ApoE4 increases total Aβ levels and the fraction of synapses containing Aβ

Aβ is present at synaptic terminals, particularly at 25 days *in vitro* (**Figure 2**). We wondered whether the extent of Aβ at synapses is also dependent on the ApoE genotype. We observed that human Aβ is colocalized to synaptic marker synaptophysin in APP^NL-F^-KI neurons in the presence of both human ApoE3 and ApoE4 (**Figure 6A**). Note that even endogenous mouse Aβ in wild-type neurons seems to colocalize with synaptophysin (**Supplementary Figure 4**) and is thus also localized to synapses. As the scope of this study was to examine the effect of ApoE isoforms on human Aβ, all further experiments were done in APP^NL-F^-KI primary neurons, where mouse Aβ is replaced by its human equivalent.

**Figure 6:**
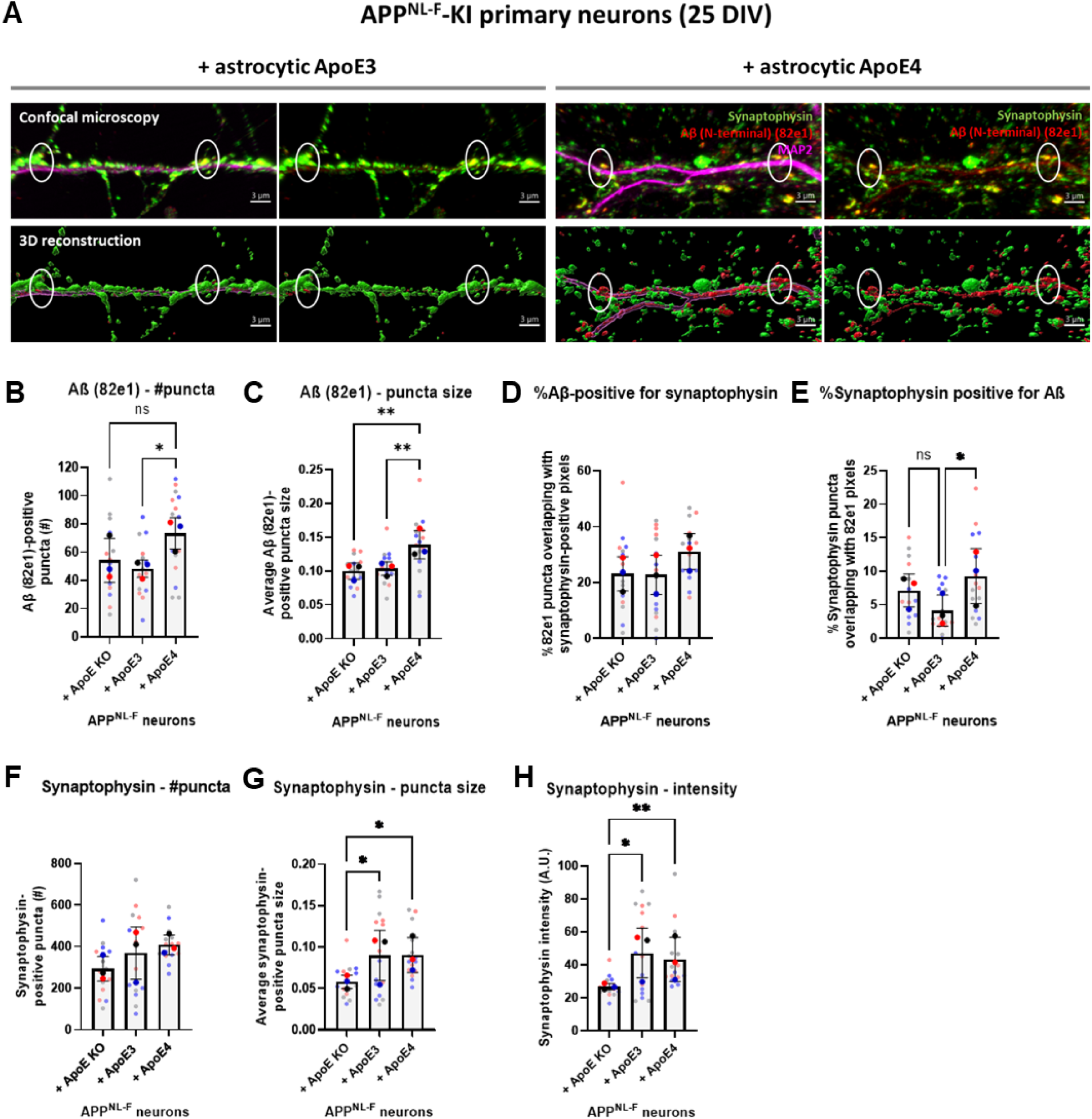
Aβ levels in APP^NL-F^ neurons increases in an ApoE-isoform dependent manner. **A.** Representative confocal (upper panel) and 3D reconstructed (lower panel) images of aged APP^NL-F^ primary neurons (25 DIV) treated with astrocyte-derived ApoE3 and ApoE4 as described in Figure 5a. These images show that Aβ (labeled with antibody 82e1 in red) colocalizes with pre-synaptic marker synaptophysin (green) in ApoE3- and ApoE4-treated APP^NL-F^ primary neurons. The Aβ-synaptophysin colocalization is highlighted by white circles. Neurites were labeled with MAP2 (magenta). Scale bar is 3 µm. **B – H.** Quantification of the number (**B**) and size (**C**) of 82e1-labeled Aβ puncta in ApoE KO-, ApoE3- and ApoE4-treated APP^NL-F^ primary neurons. **D – E.** Quantification of the percentage of Aβ-positive pixels overlapping with synaptophysin labeling (**D**) and the percentage of synaptophysin that is also positive for Aβ labeling (**E**). **F-H.** Quantification of the number (**F**), puncta size (**G**) and intensity of synaptophysin-labeled puncta (**H**) in ApoE KO-, ApoE3- and ApoE4-treated older APP^NL-F^ primary neurons. The larger data points in the graph reflect the embryos (N = 3), the smaller data points are showing the individual neurons that were analyzed (n = 15). Statistics were performed on a neuron level (**C – H**). ns = not significant, *p ≤ 0.05, **p ≤ 0.01, ***p ≤ 0.001, ****p ≤ 0.0001. Data is shown as mean ± SD.

Addition of human ApoE4 to aged APP^NL-F^-KI neurons induced a strong impact on the overall Aβ levels along dendrites compared to ApoE3. More specifically, the ApoE4 presence in APP^NL-F^-KI neurons induced a 51.6% increase in the number of Aβ puncta compared to neurons treated with ApoE3 (**Figure 6B**, p = 0.0269). Moreover, the Aβ puncta size increased by 34.1% following ApoE4 addition compared to ApoE3 (**Figure 6C**, p = 0.0079). The changes in Aβ puncta appeared to be an ApoE-specific gain of function effect, as the Aβ puncta size in ApoE4-treated neurons was also increased compared to APP^NL-F^-KI neurons treated with ApoE KO media (**Figure 6C**, p = 0.0033). While the total Aβ is increased by ApoE4, the percentage of Aβ that is overlapping with pixels labeled for synaptic marker synaptophysin were not significantly different between the different astrocyte conditioned medium conditions in APP^NL-F^-KI primary neurons (**Figure 6D**, p = 0.1536). Interestingly, the percentage of synaptophysin-positive puncta that colocalize with Aβ labeling was increased in ApoE4-treated APP^NL-F^-KI neurons compared to ApoE3-treated neurons (**Figure 6E**, p = 0.0185). More specifically, ApoE4 treatment increased the percentage of synaptophysin that was also positive for Aβ from 15.53% in ApoE3-treated neurons to 28.67% in ApoE4-treated APP^NL-F^-KI neurons. ApoE KO medium treatment did not significantly alter the ratio of synaptophysin positive puncta with Aβ in APP^NL-F^ neurons compared to ApoE3 nor ApoE4 treatment (**Figure 6E**, ApoE KO vs ApoE3: p = 0.1600; ApoE KO vs ApoE4: p > 0.9999).

To test whether the ApoE4-induced alterations in Aβ levels also change synaptic terminals, the synaptic puncta labeled with synaptophysin were further studied. The number of synaptophysin-positive puncta appear to not be significantly altered after ApoE KO, ApoE3 or ApoE4 media treatment in APP^NL-F^-KI neurons (**Figure 6F**). However, addition of human ApoE increased the size and intensity of the synaptophysin puncta (**Figure 6G-H**), as both human ApoE3 and ApoE4 treatment, compared to ApoE KO treatment induced synaptophysin puncta enlargement (**Figure 6G**, p = 0.0429 and p = 0.0384, respectively) and resulted in brighter synaptophysin labeling (**Figure 6H**, p = 0.0416 and 0.0070, respectively). No difference in synaptophysin puncta size (p = 0.9988) and intensity (p > 0.9999) were found between ApoE3 and ApoE4 treatment, suggesting that the effect on synaptic puncta in APP^NL-F^-KI neurons was an ApoE- but not ApoE-isoform dependent effect.

## Discussion

For many years, Aβ aggregation has been central to understanding the origins and progression of AD. Traditional cellular studies have relied primarily on APP-overexpressing transgenic models, which produce supraphysiological levels of APP and Aβ.^54^ Here, using primary neurons expressing human APP at more physiological levels,^22^ we reveal that the earliest steps of the pathogenic cascade involve changes in both Aβ quantity—intraneuronal and synaptic accumulation—and Aβ quality—structure. Endogenous Aβ was detectable in neurons and became enriched at synaptic terminals of APP^NL-F^ neurons as cultures aged. β-sheet–rich structures, characteristic of misfolded and aggregated proteins, were detected only in neurons expressing human mutant APP and not in neurons expressing the mouse protein, supporting that AD-linked human APP mutations are sufficient to induce structural misfolding even at physiological expression levels.

Aging, the strongest risk factor for AD, was modelled by prolonged time in culture and significantly intensified both Aβ quantity and quality changes. Neurons maintained for 25 DIV exhibited greater synaptic Aβ accumulation and more pronounced β-sheet signatures than those at 15 DIV. These findings parallel age-dependent intraneuronal Aβ increases in AD mouse models and human brains and extend earlier studies showing that β-sheet formation can occur within neurons.^42,55,56^ Our data also reinforce results from Burrinha et al. (2021), who described increased Aβ42 along neurites; here, we demonstrate that synaptic terminals accumulate Aβ. Together, these data support a model in which aging enhances both the amount of Aβ retained in neurons and the likelihood of its conformational transition into amyloidogenic structures.

Despite the age-dependent increase in synaptic Aβ accumulation, we did not observe synapse loss in APP^NL-F^ neurons. Aβ aggregates have previously been reported to be synaptotoxic in culture and *in-vivo* models.^7,9,13,15,57^ However, this might be driven by high Aβ concentration as smaller amounts of Aβ have been associated with increased neuronal excitability.^58,59^ However, not all studies have reported synaptogenic effects with picomolar Aβ, where picomolar concentrations of oligomers triggered synaptic degeneration.^60^ For example, introduction of the Swedish APP mutation into human embryonic stem cell–derived neurons, while maintaining physiological APP expression, led to altered synaptic markers, increased expression of synapse-related genes, and functional changes consistent with early synaptic dysregulation rather than overt neurodegeneration, supporting an Aβ-dependent modulation of synaptic function.^61^ In contrast to these human neuron models, *in vivo* mouse studies report synaptic deficits that strongly correlate with the accumulation of oligomeric Aβ.^13^ The discrepancy may reflect differences in Aβ aggregates species: picomolar of oligomers reflects a high level of aggregates and induce synaptotoxic effects, whereas picomolar levels of Aβ with mixed conformations can be synaptogenic. *In-vivo* AD animal studies have observed synaptotoxic effects from Aβ, mainly in the vicinity of plaques.^13^ Current evidence supports a scenario in which physiological levels of Aβ monomers and oligomers enhance synaptogenesis and neuronal excitability, but over time and with higher concentrations of Aβ aggregates, the effects shift toward synaptic hypo-excitability and synaptic degeneration.^61^ Therefore, using models that maintain physiological expression of APP is critical for studying the early events in AD pathogenesis.

ApoE genotype further modulated these early events. Neurons exposed to ApoE-KO astrocyte media displayed robust β-sheet formation, confirming that endogenous human APP^NL-F^ is sufficient to induce structural misfolding once Aβ accumulates. Interestingly, treatment with ApoE3 or ApoE4 reduced β-sheet content relative to ApoE-KO, despite increasing overall and synaptic Aβ levels—particularly in the case of ApoE4. This divergence between Aβ quantity (increased accumulation) and quality (reduced β-sheet enrichment) suggests that human ApoE isoforms influence the stage at which Aβ resides in the aggregation pathway, potentially stabilizing more soluble or oligomeric species rather than promoting fibrillar β-sheet structures in the presence of ApoE4. These findings are consistent with *in vivo* data showing that ApoE4 promotes oligomer accumulation in the brain and at synapses.^14,62^ The similar synaptic morphological changes induced by ApoE3 and ApoE4 point to a general human ApoE effect on synaptic structure that may facilitate Aβ retention, regardless of isoform.

A key limitation of microspectroscopic methods such as OPTIR is the inability to assign β-sheet structures to specific proteins which hinder spectral interpretability. However, by pairing OPTIR with immunofluorescence, we observed parallel changes in β-sheet signatures and Aβ labeling, strongly implicating Aβ as the principal contributor to the structural alterations. However, definitive proof will require correlative OPTIR and fluorescence imaging at single-molecular resolution, for example by combining nano resolution techniques such as stimulated emission depletion microscopy (STED) and scanning near-field optical microscopy (SNOM).^56^

Taken together, because synaptic terminals emerge as one of the primary sites where Aβ first accumulates and later undergoes structural transformation, the approaches demonstrated here may help uncover the molecular origins of AD and inform therapeutic strategies targeting the earliest stages of Aβ pathology. Our findings identify early alterations in both Aβ quantity (intraneuronal and synaptic accumulation) and Aβ structural alteration (β-sheet content) as the intraneuronal step in the pathogenic cascade of AD. These events are modulated by aging and ApoE genotype, two major AD risk factors. This provides further evidence that early amyloidogenic processes can arise without APP overexpression, highlighting the APP^NL-F^ model as a powerful and more physiologically relevant system for dissecting the earliest molecular events in Alzheimer’s disease.

## Methods

### List of primary antibodies used in the study

List of primary antibodies used in the study.
| Antibody | Company | Catalogue number | Dilution | Application |
| --- | --- | --- | --- | --- |
| APP (6E10) | Biologend | 803001 | 1:1000 | IF |
| N-terminal Aβ (82e1) | IBL | 10323 | 1:500 | IF |
| β-Actin | Sigma | A3854 | 1:20000 | WB |
| Bassoon | Novus Biologicals | NB120-13249 | 1:500 | WB and IF |
| Drebrin | Invitrogen | PA1-20222 | 1:1000 | WB |
| MAP2 | Invitrogen | AB92434 | 1:1000 | IF |
| PSD95 | Millipore | AB9708 | 1:200 | WB |
| Rab3a | Cell signaling tech | 3930 | 1:1000 | WB |
| Synaptophysin | Invitrogen | MA5-14532 | 1:1000 | WB and IF |

### Animals

To study human APP and Aβ the APP^NL-F^ knock-in (KI) (App^tm2.1Tcs^/App^tm2.1Tcs^) mice were used.^22^Human ApoE was studied using the ApoE KO (B6.129P2-Apoe/J; Jackson Laboratory), humanized ApoE3-KI (B6.Cg-Apoeem2(APOE*)Adiuj/J; Jackson Laboratory), humanized ApoE4-KI (B6(SJL)-Apoetm1.1(APOE*4)Adiuj/J; Jackson Laboratory) mouse models. All performed animal experiments were approved by the Ethical Committee for animal research at Lund University (ethical permits: Dnr. 5.8.18-05983/2019 & 5.8.18-13038/2024).

### Primary neuron cultures

The localization and aggregation of human APP and Aβ were studied in primary mouse neurons. Primary neurons were obtained from cortical and hippocampal brain tissue from APP^NL-F^ KI embryos (E15-17). The brain tissue was dissected and place in cold Hank’s balanced salt solution (SH30588.1, Thermo Fisher Scientific) supplemented with 0.045% glucose, followed by manual dissociation using 0.25% Trypsin (15090046; Thermo Fisher Scientific), as previously described^63^. The dissociated cells were seeded on poly-D-lysine-coated coverslips or 6 well plates in DMEM media supplemented with 10% fetal bovine serum (FBS) (10082147; Gibco) and 1% antibiotics (Penicillin/Streptamycin (p/s); SV30010; Thermo Fisher Scientific) After 4-14 hours, DMEM media was replaced by neurobasal media containing B27 supplement (17504044; Gibco), 1% p/s, and 1.4 mM L-glutamine (25030081; Gibco). Primary neurons were cultured for 15 or 25 days *in vitro* (DIV) before collection/fixation.

### Primary astrocyte cultures

Human ApoE was obtained from ApoE3-KI and ApoE4-KI primary astrocyte cultures. Primary astrocyte cultures were derived from cortical and hippocampal brain tissue from mouse pups (P1-P3) as previously described.^36^ The tissue was dissected and dissociated in 0.25% Trypsin in a manual manner using plastic Pasteur pipets. The cells were seeded onto T75 plates and cultured in AstroMACS medium (130-117-031; Miltenyi Biotec) supplemented with 0.5 mM L-glutamine. Every 2-3 days, the media was replaced by fresh media.

To collect human ApoE, conditioned media was collected from the primary astrocyte cultures once they reached at least 80% confluence. First, the astrocyte cultures were shortly washed in PBS and the AstroMACS media was replaced by neurobasal media containing B27 supplement (17504044; Gibco), 1% p/s, and 1.4 mM L-glutamine (25030081; Gibco) for 48 h. After that, conditioned media were collected on ice, centrifuged for 10 min at 10,000 x g and stored at -80°C in small aliquots until further use. ApoE KO primary astrocyte cultures were used as a control for no ApoE. When supplying primary neurons, 50% of the neurobasal media was removed and supplemented with an equal volume of conditioned ApoE astrocyte media.

### Infrared spectroscopy

Optical Photothermal Infrared (OPTIR) spectromicroscopy provides submicron spatial resolution for structural characterization of biological samples.^40^ Unlike conventional infrared modalities—such as Fourier Transform Infrared (FTIR) and Quantum Cascade Laser (QCL)–based IR microscopes—which directly measure absorbance and are therefore diffraction-limited, OPTIR detects the photothermal response generated by IR absorption, enabling sub-cellular structural measurements. The system operates using a pump–probe optical spectroscopy design. A pulsed, tunable QCL serves as the IR pump beam, inducing local photothermal heating upon absorption by the sample. This heating causes rapid mechanical and optical changes, including transient surface expansion and refractive index modulation. A co-aligned 532 nm continuous-wave green laser acts as the probe beam, detecting these photothermal changes. Modulation of the probe beam intensity at the QCL pulse frequency is isolated using a lock-in amplifier, producing high-quality infrared spectra directly comparable to FTIR while overcoming the diffraction limitations of traditional IR systems. Sample preparation for spectroscopy: neurons were grown on glass coverslips for 15 and 25 DIV. Coverslips with neurons were washed with PBS at and fixed for 15 min with 4% paraformaldehyde (PFA) at room temperature. Neurons were stored in PBS at 4° C until measurements were performed. Before measurements, neurons were washed with distilled water and dried under N_2_ flow. Measurement parameters: APD detector, probe power 0.6 %, IR laser power 22%, spectral range was set 1530-1780 cm^-1^, 10 spectra average per every acquired spectra.

### Spectroscopic data analysis

The analysis of OPTIR spectra was conducted utilizing the built-in PTIR Studio software (Photothermal Corp. St. Barabara, US). Following automatic background subtraction, the spectra were normalized to the amplitude intensity value at 1656 cm^-1^ and then averaged. To enhance spectral features and identify peak positions, the second derivative of the spectra was calculated to the second order. This process was achieved using the Savitsky-Golay algorithm with a ten-point filter and a polynomial order of three. Spectra with noticeable noise were excluded from further analysis.

For the investigation of β-sheet structures, the peak intensity ratio between 1620−1635 cm^-1^, corresponding to β-sheet structures, and 1656 cm^-1^, mainly representing cm^-1^, α-helix content, was computed. An elevation in the 1620−1635 cm^-1^ component was interpreted as a characteristic of increased amyloid fibril concentration. Spectral fitting of the OPTIR data was performed using Quasar (Orange Spectroscopy).^64^ Prior to fitting, spectra were preprocessed by baseline correction and vector normalization. To guide peak assignment, the second derivative of each spectrum was calculated, allowing identification of underlying component peaks within the Amide I region. The Amide I band (1600–1700 cm⁻¹) was then modeled using a combination of Gaussian and Lorentzian components corresponding to distinct secondary structure elements, including β-sheets (major components), α-helices/turns, and antiparallel β-sheets (minor components). Quasar’s non-linear least-squares fitting algorithm was used to optimize peak positions, widths, and intensities. The same fitting parameters, constraints, and preprocessing steps were applied to all spectra to ensure consistency across samples.

### Western blot

Synaptic proteins and ApoE were detected and quantified using western blot. Primary neurons were collected and lysed in cold NP40 buffer, consisting of 20 mM Tris, 150 mM KCl, 5 mM MgCl2, and 1% NP40 with Halt^TM^ Protease inhibitor cocktail (Thermo Fisher Scientific, 78430) and Halt^TM^ Phosphatase Inhibitor Cocktail (Thermo Fisher Scientific, 78428). The cell lysates were centrifuged at 20,000 x g for 20 min at 4°C, where after the supernatant was collected for further analysis.

To perform SDS PAGE, the cell lysates were mixed with Novex NuPAGE^TM^ LDS sample buffer (Invitrogen, NP0007) according to manufacture guidelines. 0.8% β-mercapto-ethanol was added to the LDS sample buffer to create reducing conditions. The lysates were heated at 70°C for 10 min prior to loading of the samples on a NuPAGE^TM^ 4-12% Bis-Tris protein gel (Invitrogen, NP0321). The gel was run at 150 V. SeeBlue® Plus2 pre-stained protein standard (Invitrogen, LC5925) was run at a separate lane to identify the correct molecular weight bands of the proteins of interest. After the gel electrophoresis, the proteins on the membranes were transferred to PDVF membranes using the iBlot^TM^ 2 Gel Transfer Device (Thermo Fisher Scientific, IB21001). Unspecific signal on the membrane was blocked by incubating the membrane in PBS containing 0.1% Tween (PBS-T) and 5% bovine serum albumin (BSA) for at least 1 h.

Prior to antibody incubation, the membranes were cut at different molecular weights (based on the molecular weight ladder) to allow us to study several synaptic proteins at the same protein membrane. The membranes were incubated in primary antibody incubation overnight at 4°C. Thereafter, the membranes were incubated in Horse radish peroxidase (HRP)-conjugated secondary antibodies for 1 h at room temperature. All washes performed in between the antibody incubation steps were performed in PBS-T. The antibodies were diluted in PBS-T containing 5% BSA. After the final washing step, the membranes were incubated in ECL detection reagent (170-5061, BioRad) and imaged using ChemiDoc XRS+ with ImageLab. The synaptic protein levels were quantified using ImageJ 1.53q software and were normalized to β-actin.

### Immunofluorescence

Mouse wild-type and APP^NL-F^-KI primary neurons grown on coverslips were fixed in PBS containing 4% paraformaldehyde (PFA) for 15 min at room temperature after culturing them for 15 days *in vitro* (DIV) (mature) or 25 DIV (older or aged). The fixed neurons were subsequently incubated in permeabilization and blocking solution containing of 0.1% saponin (Sigma-Aldrich, 84510), 1% BSA and 2% normal goat serum (NGS) (Jackson ImmunoResearch, 005-000-121) in PBS for 1 h at room temperature. To label our proteins of interested, the primary neurons were incubated in primary antibodies specific for synaptic proteins, APP/Aβ or a neuronal marker overnight at 4°C. The next day, the neurons were further incubated in fluorescent Alexa-conjugated secondary antibodies for 1 h at room temperature. After the start of the secondary antibody incubation, all steps were performed in the dark. All washing steps were performing in PBS. Primary and secondary antibodies were diluted in 2% NGS in PBS.

### Fluorescence microscopy

The fluorescently labeled neurons on coverslips were imaged using two different kinds of microscope. For quantification, images were taken using a Nikon Eclipse 80i upright epifluorescence microscope. The microscope was equipped with a 10x, 20x, 40 x and 60x objectives. The images in this study were all taken with the 60x (Nikon apo VC, NA 1.40, oil immersion) objective. The epifluorescence Nikon microscope was connected to a computer with Nikon Instruments Software-Elements Advance Research (NIS-Elements).

Higher resolution images and images needed to create 3D reconstruction were taken by using a laser scanning Leica TCS SP8 microscope (Leica Microsystems). The confocal Leica microscope was connected to a computer with Leica Application Suite X software. For images stacks, a z-step size of 0.5 µm was used. 3D reconstructions of our images were created using Bitplane Imaris software using the filament tool. The filament settings were set manually to avoid false positive signals in our 3D models, and the settings were kept constant among all different conditions.

### Microscopic image analysis

To analyze synaptic puncta (synaptophysin-positive and bassoon-positive), analyze particles in ImageJ software was used. First the images were pre-processed using Gaussian blur to increase the contrast between the antibody-positive signal and background noise. After that, the images were thresholded and analyzed using the Analyze particles tool. 10 non-overlapping regions of interests along MAP2-positive neurites were randomly selected within each analyzed neuron. The area of each individual region of interest was kept constant. The values of the 10 region of interest were averaged into one value for each neuron (which was shown in the graphs). The values for puncta size and number of puncta were directly obtained by Analyze particles. The intensity of the puncta was obtain by measuring the intensity (mean gray value) in the original, unprocessed image. The synaptophysin-positive puncta size was further analyzed by subdividing the puncta in three size-groups: small, medium and large. To do this, all synaptophysin-positive puncta of all groups were pooled and the mean and quartiles were determined. All puncta < first quartile were subdivided into the ‘small group’. All puncta > first quartile and < third quartile were subdivided into the ‘medium’ group and all puncta > third quartile were considered ‘large’.

Aβ-positive puncta (labeled for antibody 82e1) were analyzed using ImageJ software. Images were background subtracted and thresholded prior to analysis. Like for synaptic puncta, 10 non-overlapping regions of interest along MAP2-positive neurites were randomly selected for analysis. The region of interests was set based on MAP2 labeling to avoid bias. However, as big, potentially unspecific, clumps of 82e1 antibody sometimes appear on the coverslips and this might impact the result, the final regions of interest were checked for big, potentially unspecific, 82e1 signal before usage. The size and number of puncta was analyzed using Analyze particles, the fluorescence intensity was obtained by measuring the mean gray value of the puncta in our regions of interest in the original, unprocessed image.

The colocalization of the synaptic marker synaptophysin with Aβ (82E1 antibody) was quantified in ImageJ by calculating the percentage of overlapping pixels. Images were preprocessed as described above for the Analyze Particles workflow. After preprocessing, selections were created to include all synaptophysin-positive puncta or all Aβ-positive puncta within the defined region of interest.

Colocalization was assessed by two complementary ways: (1) Percentage of Aβ pixels overlapping with synaptophysin — representing the fraction of total Aβ located at synaptophysin-positive synapses. (2) Percentage of synaptophysin pixels overlapping with Aβ — representing the fraction of synaptic puncta containing Aβ. For each analysis, the selection (Aβ for situation 1, synaptophysin for situation 2) was overlaid with the thresholded image of the other marker. Colocalization values were obtained by measuring the area fraction of overlapping pixels within the selection and multiplying the resulting fraction by 100 to yield a percentage.

### Scanning electron microscopy

The fixed and washed neurons were critical-point dried using a Leica CPD300 system, mounted on SEM stubs, and sputter-coated with gold (Cressington 108 Auto; 55 seconds at 20 mA). The preparations were imaged with a Hitachi SU3500 Scanning electron microscope at 5kV.

### Statistics

Statistical tests in this study, except for the data in Figure 1, were performed using Graphpad Prism software. Each dataset obtained by fluorescence microscopy of western blot was tested on normal (Gaussian) distribution of the data. The normality of the data was studied through performing normality tests (Shapiro-Wilk and Kolmogorov-Smirnov tests) and QQ plots. Mann-Whitney tests were performed to compare mature (15 DIV) to older (25 DIV) APP^NL-F^-KI primary neurons (**Figure 2**, data were not normally distributed). Wild-type and APP^NL-F^-KI primary neurons at two different ages (15 and 25 DIV) were analyzed using Two-way ANOVA tests (**Figure 3 and 4**, data were normally distributed). Either a parametric One-way ANOVA or a non-parametric Kruskal-Wallis test, depending on the normality of the data, was performed for comparing ApoE KO-, ApoE3- and ApoE4-treated APP^NL-F^-KI primary neurons (**Figure 6**). Statistical analysis on data obtained from infrared microspectroscopy was done using one-way ANOVA, followed by Tukey’s pairwise comparisons test. The data in the graphs is always displayed as mean ± SD, unless stated otherwise. When super plots were used, bigger data points display the single embryos studied (N), smaller data points reflect single neurons analyzed (n). Within each neuron, 10 randomly selected dendritic fragments were studied and averaged for each neuron. The statistical tests were performed on a neuron level (n).

Statistical comparisons in Figure 1 were done *using* OriginPro^R^ by one-way ANOVA followed by Tukey’s test, with the significance level set to 0.05, the bar shows mean ± SD. For each genotype, spectra were acquired from 20–50 individual cells, yielding an initial dataset of 20–50 spectra per condition. Spectra of insufficient quality (e.g., low signal-to-noise ratio or artifacts) were excluded prior to analysis. The remaining high-quality spectra were subjected to unsupervised clustering. For downstream analyses, only spectra belonging to clusters exhibiting defined structural features of interest (e.g., β-sheet–enriched signatures) were retained.

## Supporting information

Manuscript Konings et al

## Data availability

All data presented are included within the article. The raw data supporting the conclusions of this article will be made available by the authors, without undue reservation, to any qualified researcher.

## Supporting information

This article contains supporting information

## Acknowledgments

We acknowledge MultiPark for providing access to the infrastructure. We acknowledge Ola Gustafsson at the Microscopy Facility for bioscience at the Faculty of Science, Lund University for support with Scanning Electron Microscopy and Dr. Ferenc Borondich (SMIS, synchrotron SOLEIL, France) for support with OPTIR imaging.

## Author contributions

Writing–review & editing: S. C. K., E.N., R.T. O. K., and G. G.

Conceptualization, investigation, formal analysis, methodology; S. C. K., E.N., I.M., R.T.

supervision, resources; project administration; funding acquisition G. G. and O.K

## Funding and additional information

Swedish Research Council (Grants #2019-01125,, 2024-02266);

Hjärnfonden (Grant # FO2023-0259, and FO2024-0406, FO2025-0263);

Alzheimerfonden (Grant # AF-980901).

## Supporting information

Supplementary Figures

