## Supplementary material for "Neuronal age drives β-sheet accumulation and, together with ApoE4, enhances synaptic Aβ localization in APP^NL-F^ neurons": Manuscript Konings et al

**Supplementary Information**

Sabine C Konings*^1,2^, Emma Nyberg^2,3^, Isak Martinsson^2^, Radhika Thakore^1,4^, Gunnar K. Gouras*^2,3^, and Oxana Klementieva*^1,3,4^

Affiliations:

^1^ Medical Microspectroscopy, Department of Experimental Medical Science, Lund University, Lund, Sweden.

^2^ Experimental Dementia Research Unit, Department of Experimental Medical Science, Lund University, Lund, Sweden.

^3^ Strategic Research Area Multipark, Lund University, Lund, Sweden.

^4^ Strategic Research Area Nanolund, Lund University, Lund, Sweden.

Corresponding authors:

Sabine Konings,

Gunnar Gouras,

Oxana Klementieva,

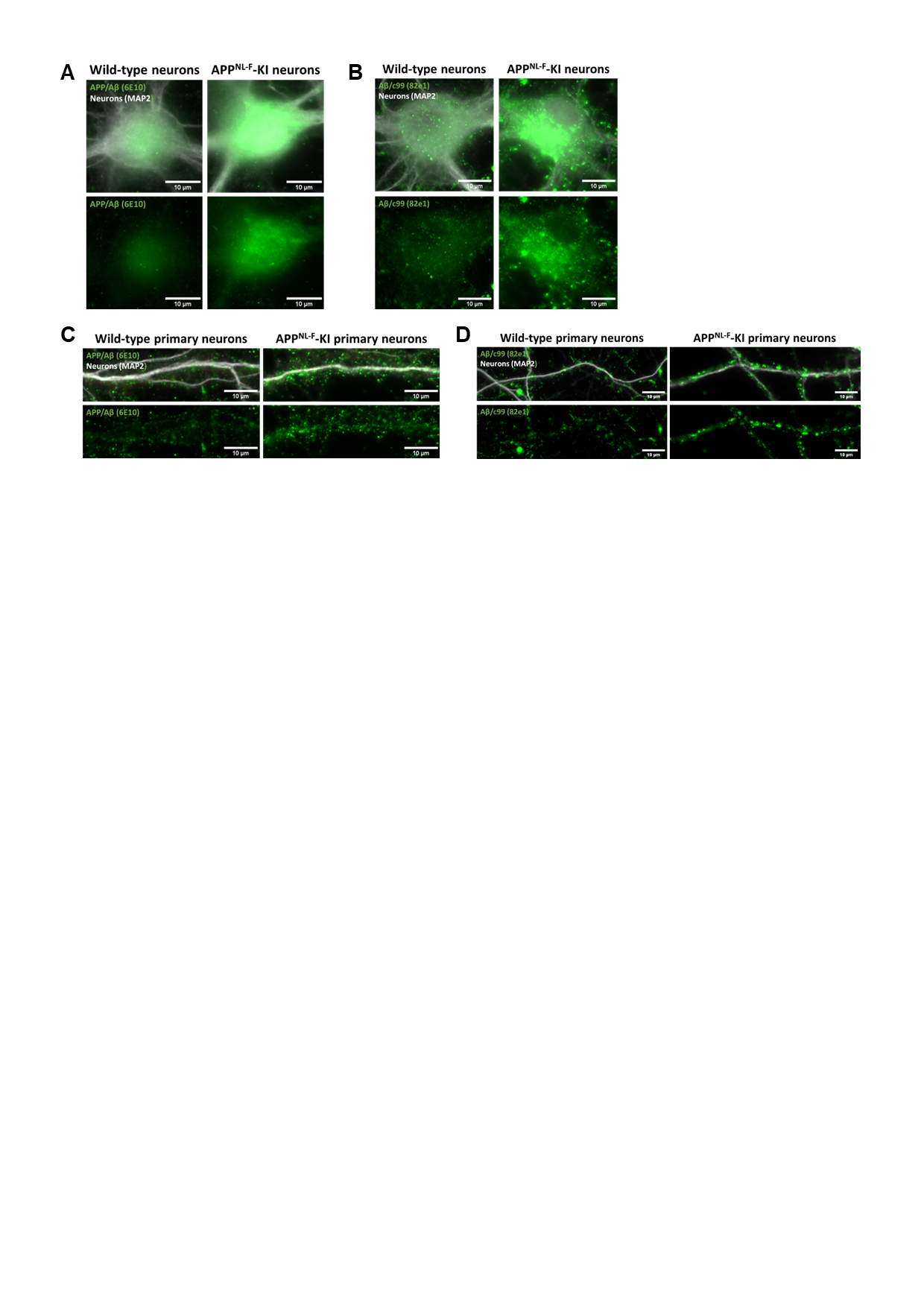

**Supplementary Figure 1: APP and Aβ immunoreactivity in wild-type and APP^NL-F^ knock-in primary neurons**. APP and Aβ are detected in mouse primary neurons derived from APP^NL-F^-KI brains. Representative images of the cell body (**A-B**) and neurites (**C-D**) of wild-type and APP^NL-F^-KI primary neurons showing 6E10 antibody-positive APP (**A, C**) and 82e1 antibody-positive Aβ (**B, D**) labeling. Scale bar is 10 µm. 6E10 and 82e1 antibody labeling is shown in green, neuronal cell bodies and neurites are labeled by MAP2 antibody, shown in white.

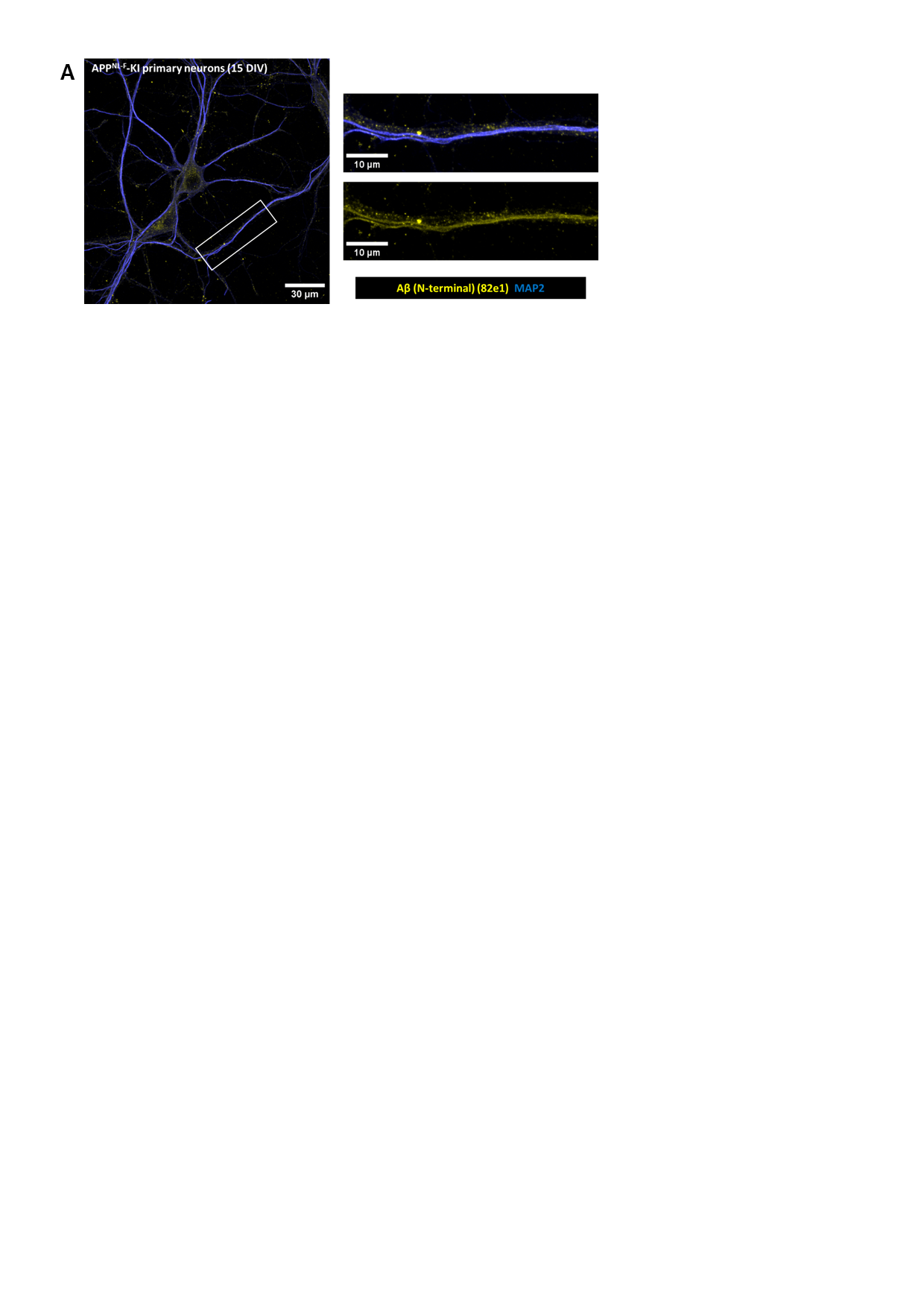

**Supplementary Figure 2: Aβ puncta in 15 div APP^NL-F^-KI neurons.** 82e1-positive Aβ labeling shows a synaptic pattern in 15 DIV APP^NL-F^-KI neurons. Representative epifluorescence images of 15 DIV APPNL-F-KI neurons labeled for Aβ antibody 82e1 (yellow) and MAP2 labeling dendrites (blue). The left panel shows an overview images of an 82e1 antibody-labeled APPNL-F primary neuron. Scale bar is 30 µm. The right panels are higher magnification images of the area indicated with a white box (left panel) showing 82e1-positive Aβ puncta along the dendrites. Scale bar is 10 µm.

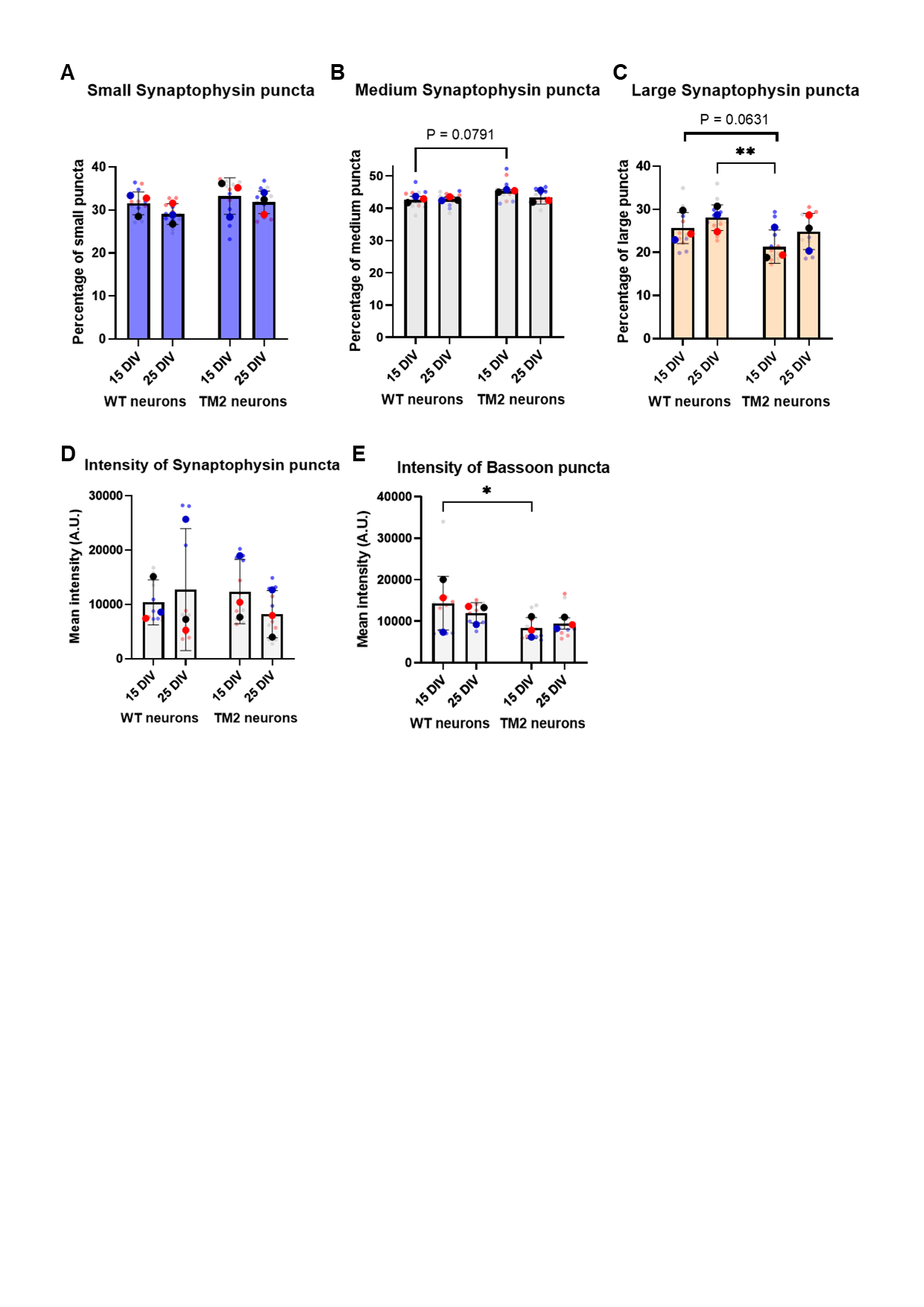

**Supplementary Figure 3:** **Changes in synaptic protein puncta**.Alterations in synaptophysin puncta size seen in APP^NL-F^ neurons are mainly caused by large synaptophysin puncta. Quantification of the percentage of small (< first quartile; **A**), medium ((> first quartile, < third quartile; **B**) and large (> third quartile; **C**) synaptophysin puncta. No alterations were detected in the intensity of synaptophysin puncta (**D**). The intensity of Bassoon puncta decreased in APPNL-F primary neurons at a mature age of 15 DIV, but not at 25 DIV (**E**).

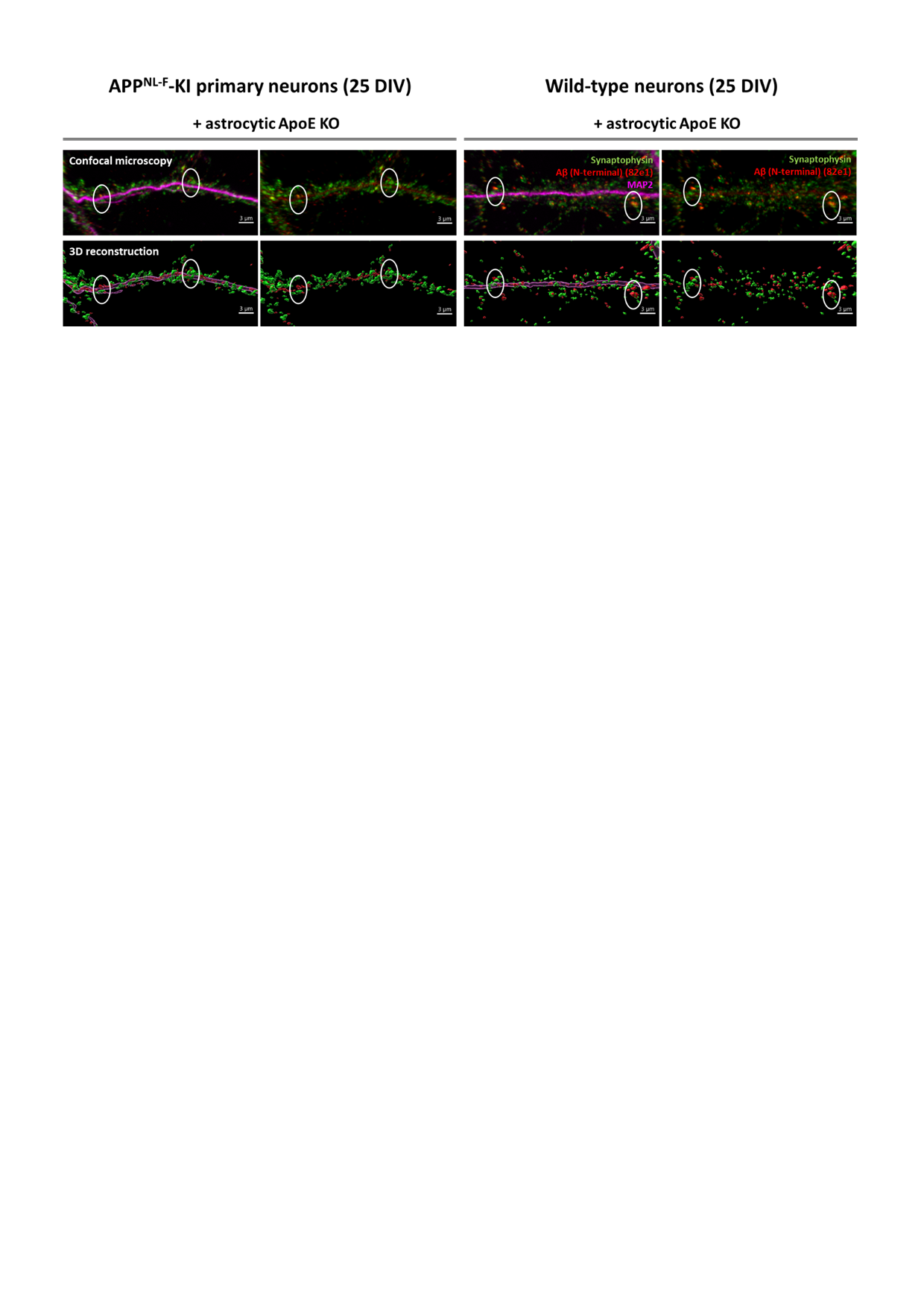

**Supplementary Figure 4: Co-localization of mouse Aβ and Synaptophysin.** Mouse Aβ in wild-type cultures neurons co-localizes to synaptophysin-positive synapses. Representative images of neurites of ApoE KO-treated APPNL-F and wild-type cultured neurons that were labeled for synaptic marker synaptophysin (green), N-terminal Aβ (82e1; red) and neuronal marker MAP2 (magenta). The white circles indicate co-localization of 82e1-positive Aβ labeling with synaptophysin puncta. These images are used as controls for the images shown in Figure 6A. Scale bar is 3 µm.

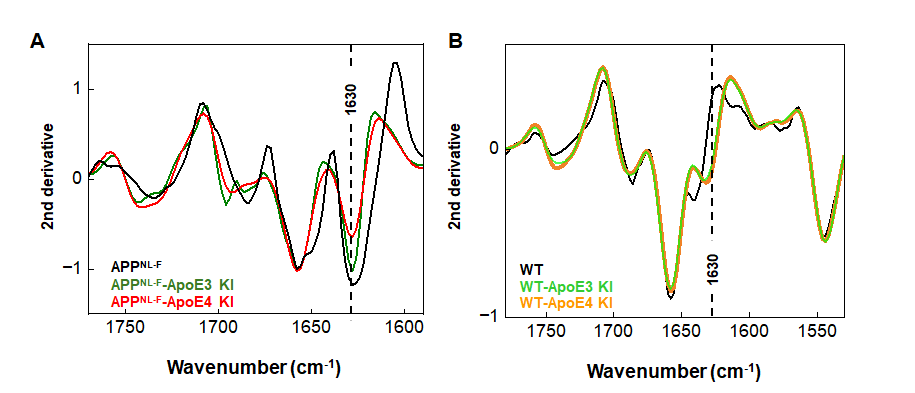
**Supplementary Figure 5: β-sheet–associated band in OPTIR spectra of primary neurons.** Averaged and normalized second-derivative OPTIR spectra of (**A**) primary APP^NL-F^ neurons incubated in astrocyte-conditioned medium and (**B**) primary wild-type neurons incubated in astrocyte-conditioned medium. Dashed lines indicate the 1630 cm⁻¹ band position.
